# Spatial statistics of submicron size clusters of activating and inhibitory Natural Killer cell receptors in the resting state regulate early time signal discrimination

**DOI:** 10.1101/2025.03.25.645117

**Authors:** Saeed Ahmad, Debangana Mukhopadhyay, Rajdeep Grewal, Ciriyam Jayaprakash, Jayajit Das

**Affiliations:** Steve and Cindy Rasmussen Institute for Genomic Medicine, Nationwide Children’s Hospital, Columbus OH; Department of Physics, The Ohio State University, Columbus OH; Department of Pediatrics, College of Medicine, The Ohio State University, Columbus OH; Pelotonia Institute for Immuno-Oncology, College of Medicine, The Ohio State University, Columbus OH; Biophysics Program, The Ohio State University, Columbus OH

## Abstract

Natural Killer (NK) cells are lymphocytes of the innate immunity and sense healthy or diseased target cells with activating and inhibitory NK cell receptor (NKR) molecules expressed on the cell surface. The protection provided by NK cells against viral infections and tumors critically depends on their ability to distinguish healthy cells from diseased target cells that express 100- fold more activating ligands. NK cell signaling and activation depend on integrating opposing signals initiated by activating and inhibitory NKRs interacting with the cognate ligands expressed on target cells. A wide range of imaging experiments have demonstrated aggregation of both activating and inhibitory NKRs in the plasma membrane on submicron scales in resting NK cells. How do these submicron size NKR clusters formed in the resting state affect signal discrimination? Using in silico mechanistic signaling modeling with information theory and published superresolution imaging data for two well-studied human NKRs, activating NKG2D and inhibitory KIR2DL1, we show that early time signal discrimination by NK cells depends on the spatial statistics of these clusters. When NKG2D and KIR2DL1 clusters are disjoint in the resting state, these clusters help NK cells to discriminate between target cells expressing low and high doses of the activating cognate ligand, whereas, when the NKR clusters fully overlap the NK cells are unable to distinguish between healthy and diseased target cells. Therefore, the spatial statistics of submicron scale clusters of activating and inhibitory NKRs at the resting state provides an additional layer of control for signal discrimination in NK cells.

**Significance:** Signal integration of opposing signals initiated by activating and inhibitory NK cell receptors (NKRs) in a noisy environment determine an NK cell’s response to healthy and diseased target cells. Superresolution microscopy imaging revealed aggregation of NKRs in submicron scales in resting NK cells. Using computational modeling, information theory, and published imaging data, we show when these clusters of the opposing NKRs are disjoint, the NK cells can separate healthy from diseased target cells but fail to do so when the clusters overlap. Thus, spatial statistics of submicron-sized NKR clusters in the resting state provide a lever for distinguishing self from non-self. The results suggest spatial organization of receptors in the resting state in may modulate signal discrimination in immune cells.

## Introduction

Natural Killer (NK) cells are lymphocytes of the innate immunity which provide important protection against viral infections and tumors(1, 2). Diverse activating and inhibitory NK cell receptor (NKR) molecules expressed on the plasma membrane of NK cells bind with cognate ligands expressed on target cells(3–6), and the integration of opposing signals within NK cells generated upon ligation of NKRs determine activating or tolerant NK cell responses(1, 2). Class I MHC molecules, present on the surface of nucleated target cells, are recognized by KIR family of inhibitory NKRs(4, 7). In contrast, ligands for activating NKRs such as NKG2D are typically expressed on tumor or virally infected cells(5). In humans, tumor or virally infected target cells increased expression of activating NKG2D ligands (referred as NKG2DL hereafter), such as ULBP1-6(8–10) and MIC-A/B(9, 11) and downregulate MHC class I expressions (12). This results in an increase in activating signals and decrease in inhibitory signals in NK cells interacting with diseased (tumor and virally infected) target cells leading to the activation of NK cells and subsequent lysis of the target cells. Healthy cells usually display null or low expressions of NKG2DL, however, NKG2DL expressions in tumor or virally infected cells can be more varied - from low to high values over two (>100 fold) orders of magnitude (13, 14).

Therefore, for an effective NK cell response to distinguish between diseased and healthy target cells, it must discriminate between target cells expressing low and high concentrations of the cognate ligands.

A wide range of imaging experiments have demonstrated spatial aggregation of several activating and inhibitory NKRs in the plasma membrane on sub-micron length scales in resting or unstimulated NK cells. Super-resolution experiments by Davis and colleagues demonstrated the existence of sub-micron sized clusters of activating NKG2D(15, 16), KIR2DS1(17) and inhibitory KIR2DL1 receptors(15, 18) in primary human NK cells and human NK cell lines in the resting state. Upon stimulation of NKRs by cognate ligands, downstream signaling reactions initiated by ligand-NKR binding can give rise to the formation of larger microscale clusters of these NKR due to cytoskeletal and/or membrane forces(19).

Similar spatial aggregation of immune receptors such as T cell receptors (TCRs)(20, 21) and B cell receptors (BCRs)(22–24) in the resting state of T- and B- cells, the lymphocytes of adaptive immunity, has been observed. These submicron scale clusters have been suggested to arise from the compartmental nature of the plasma membrane where the actin meshwork underneath the inner-leaf of the plasma membrane forms ‘picket-fences’(25, 26). The ‘picket-fence’ also acts as a diffusion barrier for the protein molecules residing within the compartments. The receptors residing in a cluster often co-localize with kinases (e.g., Lck) (25) within the same cluster or co- exist with clusters of other signaling molecules such as LAT (20), which are involved in early time signaling reactions. Thus, the formation of these clusters in the resting state could facilitate early time receptor initiated signaling reactions by increasing the local concentrations of the reactants and by bringing reactants in spatial proximity(26, 27). However, it is difficult to apply these mechanisms to NK cells as NK cell activation instead of being regulated by a single primary receptor such as TCR or BCR is regulated by integration of signals generated by activating and inhibitory NKRs. Cluster formation by activating or inhibitory NKRs can strengthen early time signaling initiated by each receptor type, however, the interaction between the activating and inhibitory signaling will depend on the spatial proximity of the NKR clusters. Recent experiments with nanofabricated patterns of NKR cognate ligands show that interaction between activating NKG2D and inhibitory KIR2DL1receptors is optimal when the separation between the receptors is about 40nm(28). Therefore, in addition to the formation of NKR clusters, NK cell signaling and activation will depend on the spatial statistics (29) of the clusters of activating and inhibitory NKRs such as the presence of any overlap between the clusters. Fluorescence correlation microscopy with NK cells in humanized transgenic mice found activating and inhibitory NKRs co-localize in submicron scale domains in the plasma membrane of hyporesponsive NK cells, whereas, the opposing NKRs reside in non-overlapping nanoscale compartments in normally responsive ‘educated’ NK cells(30). How this spatial organization of NKRs in the resting state influences signal discrimination in NK cells is not fully understood.

Here we evaluated the functional roles of submicron size clusters of activating and inhibitory NKRs present formed in the resting state in affecting early time (<1 min) NK cell signaling and ligand discrimination; to accomplish this we combined published imaging data(15, 16) within silico mechanistic signaling modeling(31), and information theory(32) for two well studied NKRs, activating NKG2D and inhibitory KIR2DL1. The relevance of spatial organization to improving reliability in biochemical signaling has been investigated in a minimal theoretical model with a single type of receptor(33). We investigated how spatial statistics of submicron scale clusters of NKG2D and KIR2DL1 influence the ability of NK cells to discriminate between low and high doses of cognate activating ligands co-expressed with inhibitory HLA-C ligands on interacting target cells. Our analysis and modeling show that submicron scale aggregation of NKG2D and KIR2DL1 in non-overlapping clusters improve the discrimination. In contrast, when the clusters of NKG2D and KIR2DL1 fully overlap, the NK cells are unable to separate signals arising from small and large concentrations of the activating ligand. Thus, formation of submicron sized clusters of activating and inhibitory NKRs in at the resting state provides an additional layer of control for signal discrimination in NK cells.

### Model

We investigated membrane proximal spatiotemporal signaling kinetics initiated by activating human NKG2D and inhibitory KIR2DL1 receptors upon binding to their cognate ligands (NKG2DL), and HLA-C, respectively. Several ligands ULBP1-6, MICA-B bind to NKG2D with dissociation constant K_D_ ≤ 1μM and unbinding rates k_off_ ∼0.02-0.0012s^-1^ (k_off_=0.023 s^-1^ (MICA)(34), k_off_=0.03 s^-1^ (ULBP1)(10), k_off_=0.022 s^-1^ (ULBP0601, an isoform of human ULBP6)(10), k_off_=0.00125 s^-1^ (ULBP0602, an isoform of human ULBP6)(10)), therefore, we used a ligand species NKG2DL with binding affinities (k_on_=2.387 x 10^-2^ *μM*^-l^*s*^-l^, k_off_=0.023 s^-1^) representing the majority of the NKG2D ligands. Since we investigate early time (≤ 1 min) signaling kinetics we considered diffusion of the NKRs and did not include any movement due to cytoskeletal forces. The model investigates spatiotemporal signaling kinetics where the NKRs, their cognate ligands, and the associated signaling proteins which are distributed in a quasi-two-dimensional simulation box representing the interface between the plasma membranes of the target cell and the NK cell and the transmembrane region in the NK cell. The simulation box is divided into 91× 91 small chambers each of volume 1_0_^2^ x*z* (1_0_ - 40*nm*) where NKRs, ligands, and associated signaling proteins react (**Figure 1A**). The height *z* of the chambers is chosen depending on the type of the molecules (details in Materials and Methods section). On average, a submicron cluster of the NKRs occupies 8 chambers. Unbound ligand molecules and signaling proteins diffuse within and outside the clusters with diffusion modeled as the hopping of molecules from one chamber to the nearest-neighbor chambers (**Figure 1B**). The NKR molecules residing within the clusters can diffuse within the cluster but are not allowed to leave the cluster. The biochemical signaling reactions considered in the model have been validated in many experiments and have been considered in computational models. The details regarding the reactions, rates, and associated references can be found in Table S1 We briefly describe the main reactions in the model here (**Figure 1A**). The binding of NKG2D with NKG2DL leads to phosphorylation of the tyrosine residues in the adaptor DAP10 associated with the transmembrane domain of NKG2D by Src family kinases (SFKs) such as Lck. Phosphorylated DAP10 recruits the guanine exchange factor Vav1, and Vav1 is phosphorylated by the SFKs.

**Figure 1.**
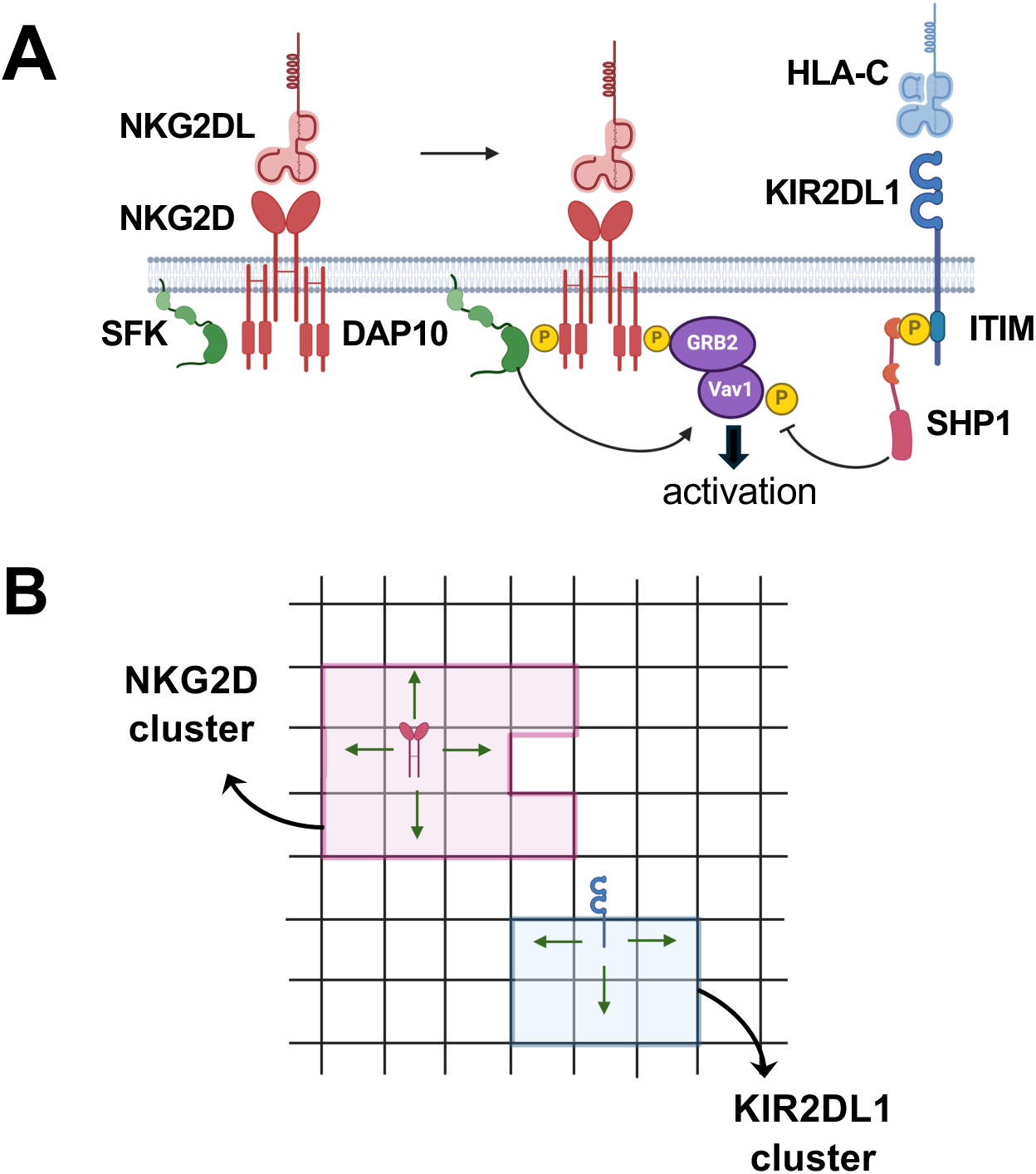
Schematic depiction of the agent- based models. **(A)** Shows the biochemical signaling reactions in the activating (leftmost) and inhibiting (rightmost) receptors used in the models. The reactions and their propensities are shown in Table S1 of the Supplementary Information Section. **(B)** Shows two examples of spatial movements considered in the agent- based models. The simulation box is divided into chambers of area *l*_0_ × *l*_0_ with a variable transverse dimension. The diffusion of receptors NKG2D and KIR2DL1 is implemented as probabilistic hops to one of their nearest-neighbor chambers, Membrane-bound molecules and cytosolic molecules diffuse at different rates, realized as different hopping probabilities to hop (see Table S1 for the parameters).

Phosphorylation of tyrosine residues on Vav1 is a key signaling event during early time NK cell signaling as phosphorylated Vav1 (or pVav1) leads to actin polymerization and degranulation in NK cells(35, 36) The inhibitory receptor KIR2DL1 binds with HLA-C, and upon binding the tyrosine in the transmembrane ITIM domains of KIR2DL1 are phosphorylated by SFKs. Fully phosphorylated ITIMs bind to the phosphatase SHP-1, and ITIM bound SHP-1 dephosphorylates pVav1 molecules that are present within length scales ≤ 40nm. The above spatiotemporal signaling kinetics in the model is simulated using kinetic Monte Carlo(37) (or spatial Gillespie simulation) which accounts for the intrinsic noise fluctuations in biochemical signaling reactions and the probabilistic nature of diffusion. Further details regarding the model are provided in the Materials and Methods section and in Table S1.

## Results

### 1. Characterization of the spatial statistics of submicron scale clusters of NKG2D and KIR2DL1 in resting NK cells

Our investigation of the effect of submicron size clusters on signaling discrimination is computed using as input the statistics of the spatial distribution obtained directly from observations reported in superresolution microscopy experiments in primary NK cells. Specifically, we characterized the spatial statistics of submicron size clusters of NKG2D(16) and KIR2DL1(17) formed in the plasma membrane of primary human NK cells in the resting state by quantifying the statistics of the spatial organization of the clusters and by the degree of overlap between the NKG2D and the KIR2DL1 clusters. First, we determined if the spatial distribution of the clusters of the NKRs observed in the super-resolution microscopy experiments exhibits any spatial order or follows a uniform random distribution. We used binary maps of the super-resolution images(16, 17) where each spatial location in the image is associated with a binary occupancy variable 0 or 1 if the site resides or does not reside within an NKR cluster. To evaluate if the NKR clusters are distributed uniformly and randomly, we generated randomized configurations of the NKR clusters where any NKR cluster is chosen randomly from the original binary image and the center of mass of the cluster in the generated configuration is chosen from a uniform random distribution. We compared the statistical similarity between the original binary images and the generated random configurations by computing the two-point correlation function C(r) for the spatial distribution of the binary occupancy variables (details in the Materials and Methods section) (**Figures 2A-B**). The C(r) computed for the original and the generated randomized images show substantial overlap (**Figures 2A-B**) revealing that the individual NKG2D or KIR2DL1 clusters in the resting state are organized uniformly and randomly in the plasma membrane. Since the imaging of NKG2D and KIR2DL1 receptors were done in separate experiments, it is unknown if the clusters of the NKRs overlap in the resting state. Therefore, we evaluated and compared the consequences of different degrees of overlap between the NKG2D and KIR2DL1 clusters for signal discrimination in our in silico study. In addition, we evaluated the effect of different organizations of the clusters on signal discrimination by comparing the results for the observed distribution of clusters with homogeneous and random distribution of NKG2D and KIR2DL1 receptors in the plasma membrane. We discuss these results in the following sections.

**Figure 2.**
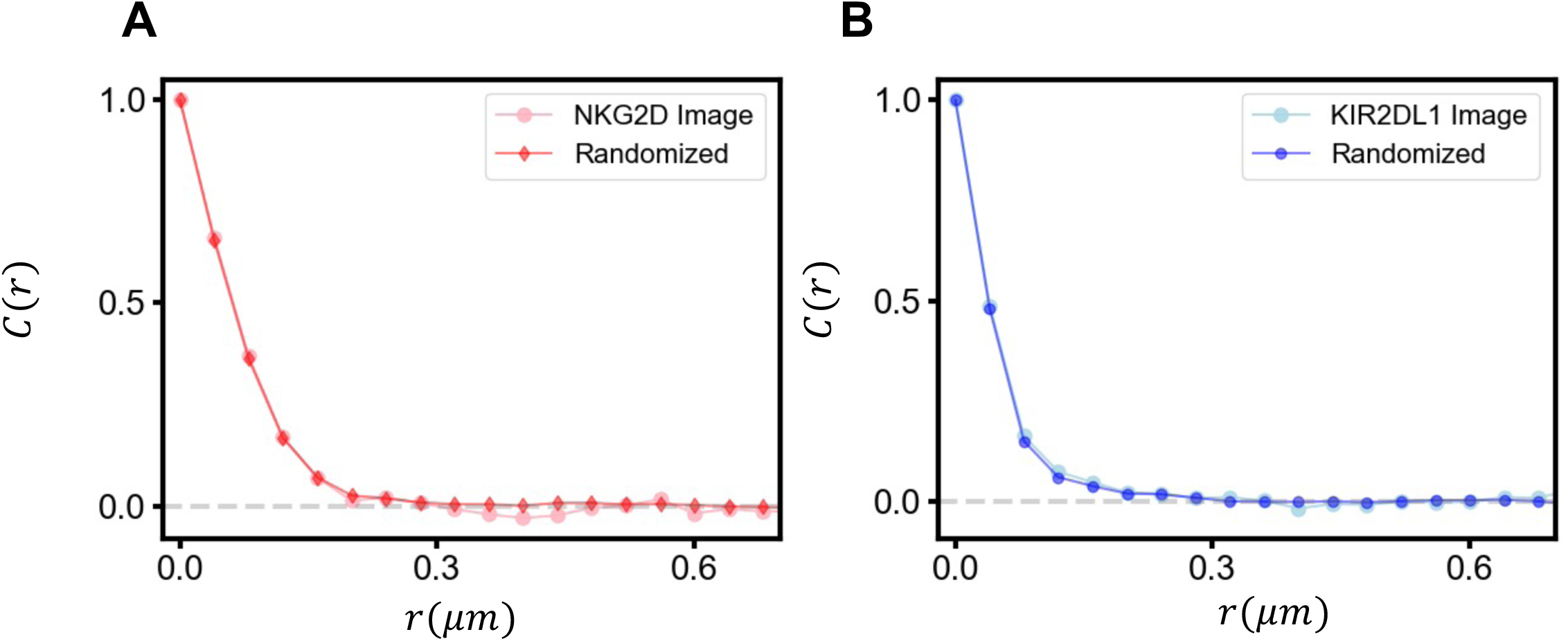
Spatial Correlation functions for superresolution images and random configurations. Superresolution microscopy images and randomly generated configuration of NKR clusters are converted into binary variables and the two-point correlations function in space are calculated as described in Materials and Methods. The substantial overlap for (A) activating NKG2D and (B) inhibitory KIR2DL1 receptors supports the conclusions (assumption?) that the clusters in the resting state are distributed uniformly and randomly.

### 2. Sub-micron clusters of NKG2D and KIR2DL1 in resting NK cells modulate early time phosphorylation of Vav1 and stochastic fluctuations of phospho-Vav1 concentrations

To evaluate the effect of overlapping submicron sized clusters of NKG2D and KIR2DL1 on NK cell response, we investigated two scenarios where the NKG2D and KIR2DL1 molecules in the resting state (i) form disjoint clusters (**Figure 3A, left**) or (ii) fully overlapped clusters (**Figure 3A, middle**). We compared results obtained for the above cases with a reference case where the NKG2D and KIR2DL1 molecules are distributed (iii) homogeneously and randomly (**Figure 3A, right**) in the simulation box. The first two scenarios allowed us to investigate the role of overlap in the clusters of NKG2D and KIR2DL1 in the resting state in influencing very early time (<1 min) signaling kinetics. This initial signaling events in turn affect later (> 1 min) signaling events as well as further clustering of the receptors and select proteins. We considered two types of target cells - healthy and diseased (virally infected or tumor) expressing normal and lower copy numbers (or abundances) of HLA-C molecules, respectively. Tumor and HCMV infected cells can express NKG2D ligands (NKG2DLs) whereas most healthy cells do not express NKG2DL under normal conditions. Tumor and HCMV infected non-tumor target cells also decrease HLA-C expressions(12), however, a sub-population of HCMV infected cells could express HLA-C abundances similar to that of healthy target cells(13).

**Figure 3.**
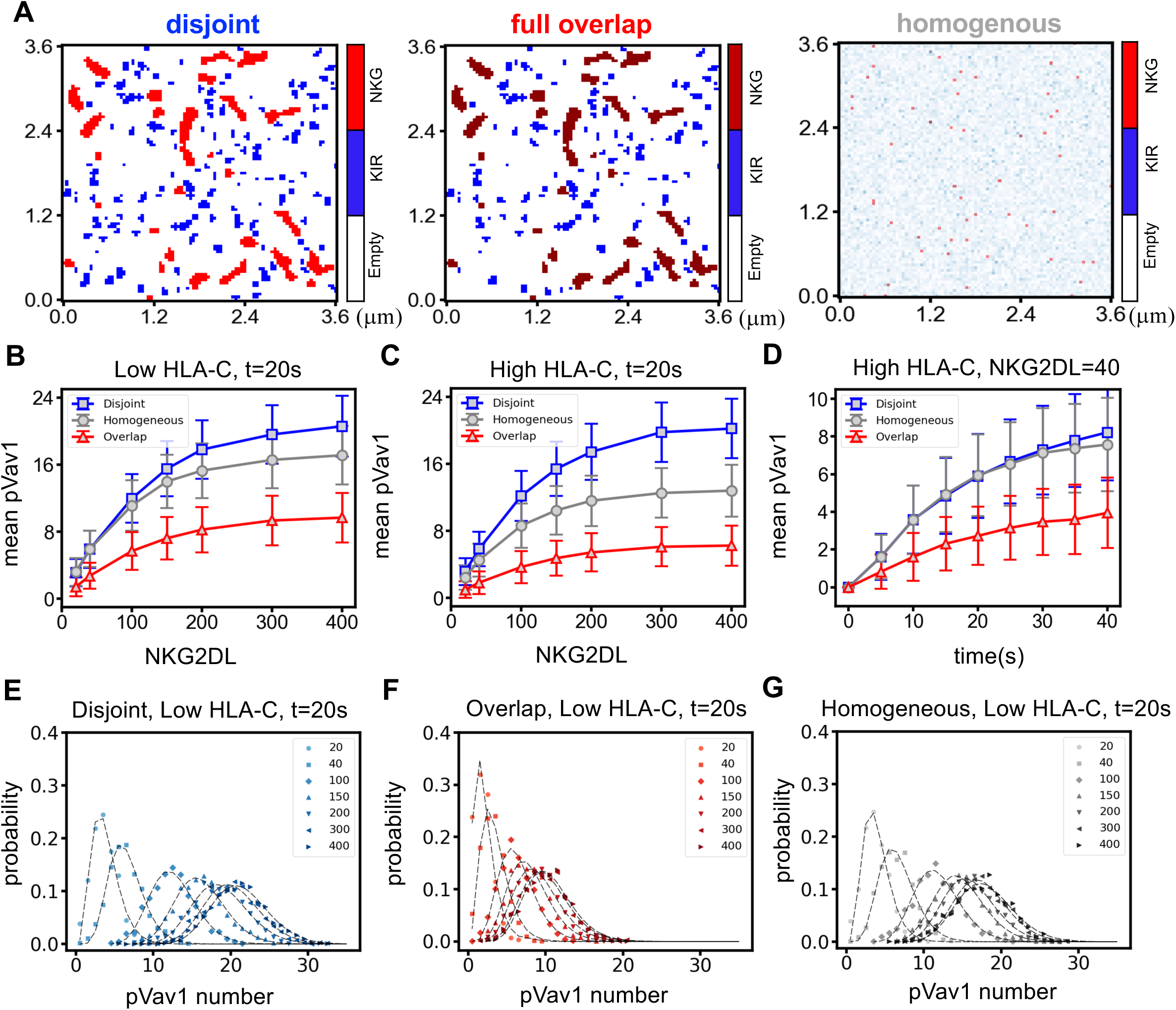
Characterizing stochastic pVav1 response for three spatial configurations as the strength of the stimulus (NKG2DL) is varied. **(A)** The three panels illustrate the spatial distribution of the NKG2D(red) and KIR2DL1(blue) receptors used in the study. The left panel shows the NKG2D and KIR2DL1 clusters distributed independently randomly with no overlap; in the middle panel the clusters overlap fully while in the third panel the two kinds of receptors are homogeneously distributed in the simulation box. **(B-C)** Shows the dependence of the mean pVav1 abundances (response) at t=20s on the copy number of NKG2DL (stimulus) when co- expressed with (B) low HLA-C (corresponding to 4213 molecules in the simulation box) and (C) high HLA-C (11686 molecules) numbers. The mean pVav1 number is substantially lower in the fully overlapping configuration for both the low and high HLAC. The response is unaffected when the HLA-C number is increased in the disjoint case in contrast to the homogeneous configuration. **(D)** Shows time dependence of pVav1 abundances for the disjoint, overlapping, and homogeneous conditions for an NKG2DL abundance of 40 molecules in the simulation box co-expressed with low numbers of HLA-C. The pVav1 abundance increases with time as more NKG2D-NKG2DL complexes are formed and reaches a steady state value that depends on the spatial configuration of the receptors, disjoint, overlapping or homogeneous. **(E-G)** pVav1 distributions for different scenarios with low HLA-C abundance and NKG2DL=20,40,100,200,400 molecules in the simulation box. The more the distributions for different NKG2DL values overlap, the less able the NK cell is to discriminate between them. The overlap is highest when the two kinds of clusters are overlapping (G) while the disjoint case(E) does slightly better than the homogeneous case (F).

To study the very early time signaling kinetics in NK cells stimulated by NKG2DL and inhibitory HLA-C ligands where the NKG2D and KIR2DL1 receptors are organized in disjoint or fully overlapping sub-micron clusters in the resting state, we simulated the spatially resolved signaling model described in Figure 1 for different copy numbers (or abundances) of the ligands NKG2DL and HLA-C per cell. The model describes the interaction between a single NK cell expressing fixed abundances of NKG2D and KIR2DL1 receptors and a target cell expressing a low or high abundance of HLA-C ligands and a specific abundance of NKG2DL. We simulated the time evolution of the system for multiple initial configurations where each initial configuration contained the same numbers of NKG2D and KIR2DL1 clusters and the same concentrations (molecules/μm^2^ or molecules/(μm^3^)) of the NKRs, ligands, and the signaling proteins, however, the spatial locations of the receptor clusters and that of the receptor and the ligand molecules in the simulation box varied between the initial configurations. The locations of the receptor clusters and the ligand molecules were chosen from uniform random distributions where specific constraints on nucleating the NKR clusters were imposed to generate disjoint and fully overlapping NKR clusters (see Materials and Methods). We used a concentration of 4 molecules/(μm)^2^ and 882 molecules/(μm)^2^ for NKG2D and KIR2DL1, respectively, which frequently generated several empty clusters devoid of NKR molecules. Given the larger concentration of KIR2DL1, empty KIR2DL1 clusters occurred rarely. The empty clusters do not contribute to the signaling kinetics.

We considered phosphorylated Vav1 (denoted henceforth as pVav1 for notational compactness) as a key marker for early time NK cell activation in our simulation as generation of pVav1 in early signaling in NK cells leads to effector functions such as mobilization of lytic granules or release of cytokines(1, 2, 38). The kinetics of the copy number of the total amount of pVav1 in the simulation box which includes unbound pVav1 molecules as well as molecules of pVav1 bound to NKG2D complexes was analyzed in detail. The intrinsic stochastic fluctuations in biochemical reactions and the probabilistic nature of the diffusion of molecules lead to the variations in the copy numbers of pVav1 in response to identical stimuli (e.g., the same abundances of NKRs and cognate ligands) in our simulations. The distribution of pVav1 copy numbers in the simulation box at any time can be viewed as the pVav1 abundances in a clonal population of NK cells where each cell contains the same abundances of NKRs and signaling molecules. We studied features of the pVav1 probability distributions generated from an ensemble of initial configurations as a function of the number of NKG2D ligands and examined their effect on the signaling kinetics. We first considered the simplest characteristic of the probability distribution, the mean pVav1 copy numbers, and investigated its time dependence and its dependence on NKG2DL and HLA-C abundances. The mean pVav1 copy numbers at t=20s (**Figures 3B and 3C** and Fig. S1A) or 40s (**Figure 3D and** Figs. S1B-C) increase monotonically as NKG2DL concentrations increase from 1.5 molecules/(μm)^2^ (=20 molecules in the simulation box) to 30 molecules/(μm)^2^ (=400 molecules in the simulation box); the mean pVav1 copy numbers are larger when the clusters are disjoint compared to those of the fully overlapping case. The homogeneous distribution of the NKRs produce mean pVav1 abundances intermediate between the disjoint and the fully overlapping case (**Figures 3B-C**, and Figs. S1A- E). This is easily understood because in the disjoint case, KIR2DL1 complexes remain spatially separated from the NKG2D-NKG2DL complexes, and are therefore unable to dephosphorylate any pVav1 bound to the NKG2D-NKG2DL complex. The negligible interaction between the NKG2D and KIR2DL1 signaling for the disjoint clusters is further validated as the simulations showed the presence of almost vanishing copy numbers of free cytosolic pVav1 molecules which can diffuse across NKG2D clusters and so potentially be dephosphorylated by SHP-1 bound to KIR2DL1. In contrast, for overlapping clusters of the NKRs, most of the pVav1 molecules encounter SHP-1 bound to KIR2DL1 complexes and can thus become dephosphorylated. When both NKRs are distributed homogeneously, about 75% of the NKG2D molecules are spatially co-localized with KIR2DL1 at t=0 due to random chance (see Supplementary Text 1). This fraction of NKG2D colocalized with KIR2DL1 does not change appreciably as the NKRs diffuse and participate in signaling reactions. Therefore, in the homogenous case, the pVav1 molecules generated by the triggering of NKG2D receptors are colocalized with KIR2DL1, can become dephosphorylated. Consequently, the mean pVav1 copy number in this case is intermediate between the mean pVav1 for the cases of disjoint and fully overlapping NKR clusters (**Figures 3B-D**, and Figs. S1A-E) for a range of NKG2DL concentrations and simulation times. When the HLA-C concentration is larger, the formation of greater numbers of HLA-C-KIR2DL1 complexes reduce the mean pVav1 copy number for the homogeneous case noticeably below that for the disjoint case (**Figure 3C**). Furthermore, an examination of the standard deviations around the mean pVav1 abundances across several NKG2DL abundances indicates overlap of pVav1 distributions for different NKG2DL concentrations (**Figures 3B and 3C**, and Figs. S1A- E). The mean pVav1 copy number for low (≈ 40) and intermediate (≥ 100) concentrations of NKG2DL showed overlap within standard errors indicating that when NKG2D and KIR2DL1 clusters in the resting state fully overlap the NK cells will be unable to discriminate between low (∼ 40) and medium (≥ 100) strength NKG2DL signals at early times (<1 min). This holds true for both low and high HLA-C abundances that are distributed homogeneously in space.

The mean pVav1 copy numbers show monotonic increase with time for low NKG2DL concentrations from 0 to 40secs (**Figure 3D** and Figs. S1D-E), whereas the mean copy number reaches saturation in about ∼20 secs at high NKG2DL concentrations (Fig. S1E). We estimated the time scales determined by the local biochemical reactions by solving the deterministic rate equations for the disjoint case (the reactions and rates are in Table I of the SI): we find that an almost steady state is reached in about 3-4 seconds and therefore, the longer time scales are determined by diffusion of NKG2DL.

Initially, the NKG2DL molecules are placed uniformly randomly in the simulation box and so the time for the binding of the NKG2DL to NKG2D molecules is controlled by how rapidly a diffusing NKG2DL molecule encounters a NKG2D next to since the binding reaction itself is rapid. There are 53 NKG2D molecules and the abundance of NKG2DL varies between 20 and 400 and the time for the diffusive encounter can be estimated as follows. Imagine tessellating the simulation box into Voronoi cells consisting of all sites closer to the ligand than to any other.

The time it takes for the NKG2DL ligand to visit the entire area of the Voronoi cell by diffusion gives an upper bound on the time for it to encounter a NKG2D. The average number of chambers of side *l*_0_ in a Voronoi cell is 8281/M where M is the number of NKG2DL molecules and there are 8281 chambers in the simulation box. Since it is known that the number of distinct sites visited in an *n*-step random walk on a square lattice is very well approximated by ∼ π*n*/Log[5.692*n*](39, 40), the average time τ to visit *n*_v_ number of sites in a Voronoi cell is as follows. For a ligand taking an average time *t*_step_ (∼ 1/6 sec for NKG2DL) to take a step of length *l*_0_ the number of sites visited in a time τ is given by, *n*_v_ ≈ π(τ/*t*_step_)/Log[5.692(τ/*t*_step_)]. Thus, for M=200, when most Voronoi cells do not contain a receptor, all the chambers will be visited by 15s and including the time for biochemical kinetics, saturation starts occurring between 15 and 20s; this is consistent with **Figure S1E**. For M=40, since there are 53 NKG2D, some of the Voronoi cells will have more than one receptor, but not all receptors are bound at a given time. In this case around half of the chambers in a Voronoi cell are visited by about 40s; This is roughly consistent with **Figure 3D**.

To examine the variations of pVav1 copy numbers for a given concentration of NKG2DL and HLA-C, we present the probability distributions of pVav1 copy numbers obtained from at least runs of the simulation for the same NKG2DL and HLA-C concentrations at t=20s (**Figures 3E- 3G**, and Figs. S1F-H) and 40s (Figs. S1I-N). The distributions of pVav1 copy numbers for low and intermediate-to-high concentration of NKG2DL clearly show greater overlap as the NKG2D and KIR2DL1 clusters in the resting state varied from the disjoint to the fully overlapping case (**Figures 3E and 3F**). When the NKRs are distributed homogeneously at t=0 (**Figure 3G**), the overlaps in the distributions of pVav1 copy numbers lie between the above cases. When there the pVav1 distributions for different stimulus strengths (NKG2DL concentrations) overlap, discrimination between those stimuli is poor. The pVav1 distributions can be fitted with Gamma distributions (**Figures 3E-G**) and using the fitting parameters we find that the KL divergence between the pVav1 distributions for NKG2DL = 400 and 20 with low HLA-C are 11.1, 2.2, and 4.7 for the disjoint, overlapping, and homogeneous cases, respectively. This is another measure of how much better the disjoint cluster configuration performs in discriminating between high and low NKG2DL concentrations (see Supplementary Text 2 for details). To further quantify the capability of the NK cells to differentiate between low and high concentrations of activating ligands we calculated mutual information and channel capacity for the signal transmission in NK cells interacting with target cells expressing a range of NKG2DL concentrations.

### 3. Information-theoretic characterization quantifies the reduction in the discrimination ability between different NKG2DL abundances due to the overlap in submicron scale clusters of NKG2D and KIR2DL1 in resting NK cells

The overlap of the distributions of pVav1 copy numbers for different concentrations of NKG2DL reduces the ability of the NK cell to discriminate between healthy and diseased target cells. We quantified the ability of NK cells to discriminate between target cells expressing low and high concentrations of NKG2DL using mutual information (MI) and channel capacity (CC). Signaling in NK cells transmits information about stimulus from the extracellular environment via the NKRs and chemical modification of signaling proteins (e.g., pVav1) regulated by NKR stimulation in the cytosol through a cascade of reactions to the nucleus of the NK cells to generate an appropriate functional response. We view the stimulus NKG2DL concentration for a fixed NKG2D concentration as the input, and the phosphorylated Vav1 whose abundances encapsulate the transmitted information within the NK cell about the stimulus as an output. The intrinsic stochastic fluctuations in biochemical reactions and the diffusion of ligands and receptors lead to variations of pVav1 abundances resulting from identical stimuli (e.g., same concentrations of NKRs and cognate ligands) which can be described by distribution of pVav1 copy numbers in a population of NK cells with the same concentrations of NKRs and signaling molecules; the overlap of the distributions of pVav1 copy numbers for different strengths of stimuli decreases the ability of the cell to generate a functional response required for that stimulus, in other words, the fidelity of information transfer. Mutual information is a measure of the fidelity of output to input signals (41) and has been used in diverse areas (42, 43) and is defined as follows. The signal (*s*), in our case the concentrations of NKG2DL expressed on the surface of target cells, is assumed to follow a distribution we denote by *Q*(*s*). The NK cells interact with the target cells resulting in the generation of pVav1 copy numbers which is the output (*o*) in our case. The distribution of pVav1 copy numbers in the NK cell population, denoted by *R*(*a*), results from the interaction of the NK cells with the target cells, and the joint probability distribution for the NKG2DL concentrations and pVav1 copy numbers is denoted by *P*(*a,s*). The mutual information (MI) is defined by:

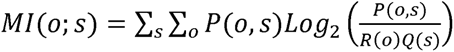

The MI is non-negative, i.e., MI≥0. Given a non-zero value of the MI, the system output can distinguish about 2*^MI^* different input signals, i.e., for MI ≈, two different input signals (e.g., low and high) can be distinguished(41). Note that the numerical value of MI depends on the form of the input signal and is not an intrinsic property of the signaling network. A quantity that is a measure of the intrinsic signal processing fidelity is the so-called channel capacity (CC), the maximal possible information that can be transduced by the system given its noise profile obtained by maximizing the mutual information over all possible input distributions: *CC* = *max_Q_*_(*s*)_MI(o;s). Thus, *CC* measures the optimal ability of the system to discriminate between input signals. We employed the Arimoto-Blahut algorithm(44, 45), an iterative algorithm that determines the value of the maximum information and the input probability distribution that leads to it. The algorithm is easily implemented and provides a computationally tractable way of determining the capacity. When the system is deterministic, i.e., a unique input signal results in a unique output signal or the conditional probability *P*(*o*|*s*)=1, the M1 is equal to Shannon entropy *H*(*s*) (see Materials and Methods) which is maximum for the uniform distribution of *s* in a fixed range. This implies that in the deterministic limit the MI value calculated with a flat signal distribution will be maximal. So mutual information calculated with a uniform input distribution, which is computationally convenient, is a good proxy for the information capacity and so we use a uniform distribution of NKG2DL concentrations to compute MI.

We computed mutual information for a uniform distribution of NKG2DL concentrations - equal probabilities for a few select values of their number between 20 and 400 molecules in the simulation box to reduce computational cost. We compute MI, as well as, the channel capacity *CC*, for an NK cell expressing NKG2D and KIR2DL1 receptors interacting with target cells expressing ligands NKG2DL and HLA-C distributed in a uniform distribution for the three types of spatial distributions of NKG2D and KIR2DL1 receptors in the resting state: (i) disjoint clusters, (ii) fully overlapped clusters, and (iii) homogeneously distributed in the plasma membrane. We computed the MI (**Figure 4A**) and the channel capacity *CC* (**Figure 4B**), for the pVav1 copy numbers at t=20s post-receptor stimulation. The MI values in the case of disjoint clusters are greater than 1 for both low and high values of the inhibitory ligand number; in contrast, the fully overlapping clusters produced MI values less than unity. The value of CC are less than unity for low and high HLA-C abundances suggesting that the fully overlapping clusters of NKG2D and KIR2DL1 are unable to clearly distinguish between even high or low abundances of NKG2DL. However, when the clusters are disjoint the NK cell can separate high and low NKG2DL concentrations in the target cell. The values of MI for the homogeneously distributed receptors decrease below unity and the CC values are lower than that of the disjoint but substantially more than the fully overlapping case (**Figures 4A and 4B**) and stayed over unity. Therefore, the formation of disjoint sub-micron clusters in the resting state aids clear discrimination between target tumor cells expressing high and low NKG2DL concentrations even when the concentration of HLA-C varies in the infected/tumor target cells, whereas the homogeneously distributed NKRs can reliably distinguish between high and low NKG2DL signals only for bimodal distribution of HLA-C concentrations.

**Figure 4.**
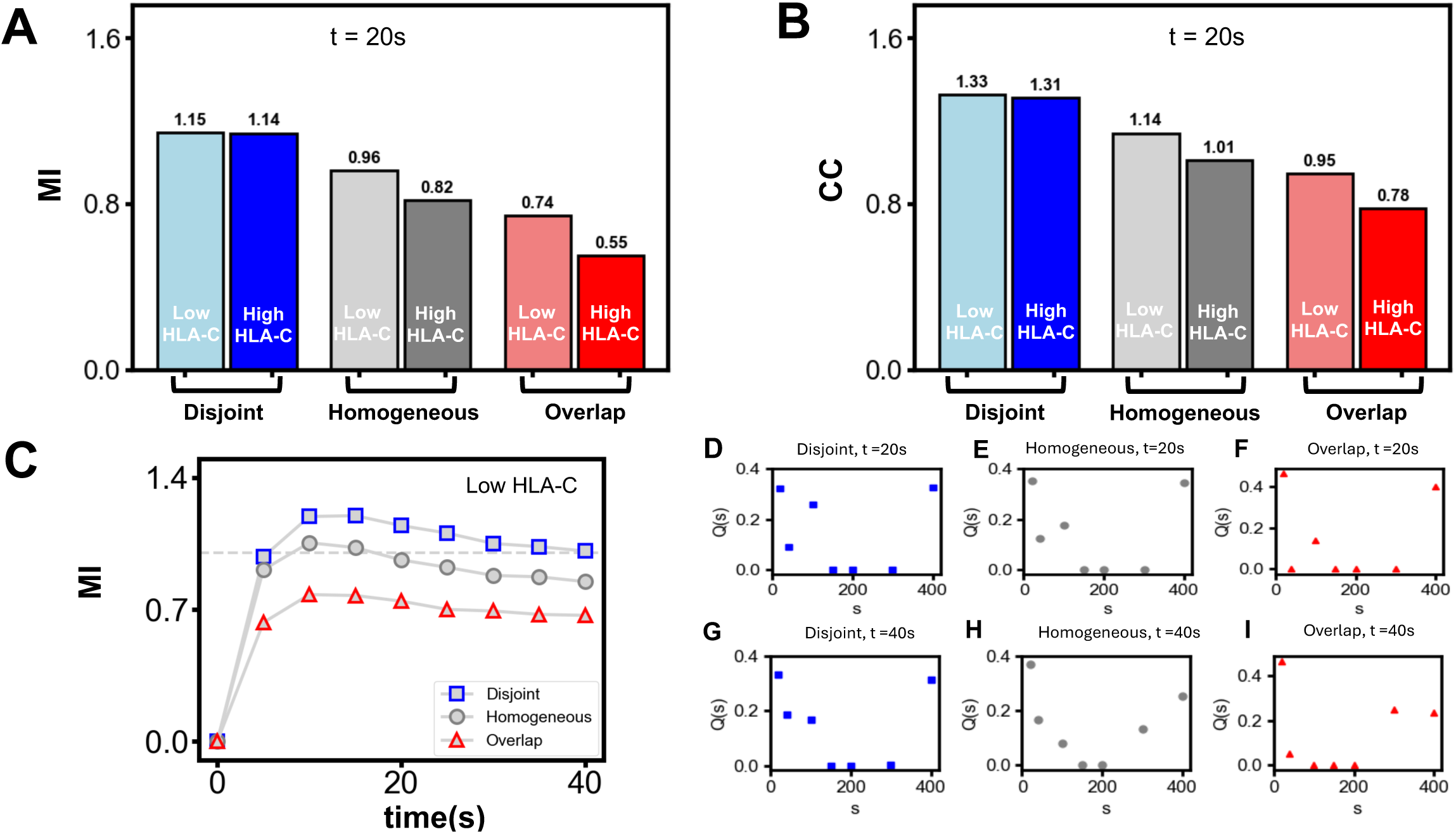
Mutual Information (MI) and Channel Capacity (CC) behavior under different conditions. **(A)** The MI values shown as bar graphs for the three spatial distributions. The dark color bars are for high HLA-C, 11686 and light color bars are for low HLA-C, 4213. Disjoint clusters exhibit the highest MI; the value is greater than 1 for both high and low HLA-C. This implies that in this case the NK cells can discriminate between two input signals (high and low) with good fidelity. The MI value is much lower for fully overlapping clusters, consistent with overlapping pVav1 distributions in Figure 3(G). In the homogeneous case MI falls from above to below unity when the HLA-C number is increased. **(B)** The channel capacity (CC), determined as the maximum MI value when the distribution of the input signal is varied, displays similar trends as the MI shown in (A). **(C)** The time evolution of MI under the three spatial conditions. It increases from zero and saturates with a slight decline beyond, e.g., after ∼ 20s in the disjoint case. When the number of bound activating receptors saturates for large NKG2DL values, their pVav1 distributions begin to coincide and the cell cannot discriminate between them though small values of input signals can still be distinguished **(D-I)** The distribution of input signals (s) that gives rise to the maximum possible mutual information denoted by Q(s) found by the Arimoto-Blahut algorithm is displayed in different cases at times 20s (D-F) and 40s(G-I), Note that it is almost bimodal with peaks at low and high signal values reflecting the two signals that the cell can distinguish.

The variation of MI (**Figure 4C** and Fig. S2A) and CC (Fig. S2B) with time for NKG2DL stimulation shows increase from small (<1) values to above unity values on time scales as short as 10 secs when the NKG2D and the KIR2DL1 in resting state are present in disjoint clusters or distributed homogeneously implying the signal discrimination in NK cells occurs in a short time scale. The MI for the overlapping case remains below unity throughout the time course (**Figure 4C**). The number of activating receptors is fixed at a value of 53 molecules in the simulation box and at early times the number of ligand-receptor pairs that are formed is diffusion-limited. The MI values increase and saturate between 10 and 15 seconds. This is consistent with the time required for the formation of NKG2DL-NKG2D complexes.

Using Mathematica (https://www.wolfram.com/mathematica/) we solved the deterministic rate equations (14 coupled ordinary differential equations for the disjoint case involving only activating receptors) that describe the local biochemical reactions once the NKG2D-NKG2L complex is formed. From the numerical solution of the deterministic rate equations, we find that for one receptor-ligand complex present initially in the simulation box, the average number of pVav1 reaches a steady-state value of slightly below 0.4 in under 5 seconds. Therefore, given a maximum of 53 NKG2D receptors we should expect a maximum of around about 20 pVav1 molecules at later times, as more and more receptors are bound to the diffusing ligands; this is consistent with the results for the time dependence of the mean pVav1 values displayed in Figure S1. The time dependence of the MI is shown in **Figure 4C** for three cases. We note that the MI is seen to decrease slightly beyond 20s. One can understand this as follows. The number of NKG2DL bound complexes increases more at early times t≤40s, then slowly saturating to a value limited by the total number of NKG2D (Fig. S4). Therefore, at later times the pVav1 distributions for 200 and 400 overlap more (**Figure 3D** vs Fig. S1I) and one would expect this to decrease the relative ability to distinguish between these NKG2DL values leading to a slight decrease in MI. However, small and large values can be separated with no reduction in fidelity and MI remains above unity.

The distributions of input signals (or NKG2DL concentration) Q(s) that maximizes MI thus yielding the CC when the NKRs in the resting state are organized in disjoint, overlapping clusters or are distributed homogeneously are shown in **Figures 4D-4I**. They are approximately bimodal distributions of NKG2DL with peaks at low and high values of the signals. Thus, when the input distribution itself is bimodal leading to CC values equal to or greater than unity (disjoint clusters or homogeneous distribution of NKRs), the NK cells would be able to separate the NKG2DL concentrations distributed in a target cell population with peaks at high and low values.

### 4. Signal discrimination across different concentrations of NKG2DL is not affected by kinetic proofreading in signaling events

Kinetic proofreading (KP)(46–50) is a mechanism has been proposed to provide improved discrimination between ligands in T cell signaling when different ligands bind to T-cell receptor (TCR) with similar affinities (K_D_ ∼ 10 – 100 μM (51); k_off_ ∼ 0.01– 0.1 s-1(51)). The presence of KP increases the specificity of discrimination by an irreversible step in the signaling kinetics that reverts an activated TCR complex to the native unstimulated state immediately once the ligand unbinds from the activated complex. KP has been shown to be present in early time TCR signaling in experiments and has been shown recently to increase the channel capacity in early TCR signaling (46). NKG2D binds to a range of ligands (MIC-A, MIC-B, ULBP1-6 in humans), however, the binding affinities of NKG2DL to NKG2D are stronger, e.g., K_D_ values ≤ 10 fold smaller than those between TCR and ligands. Moreover, the lifetimes of the NKG2DL-NKG2D complexes (∼ 20 secs(34) or longer(10)) correspond to the lifetimes of the TCR complexes formed with strong affinity ligands. Therefore, we reasoned that discrimination across different concentrations as opposed to different affinities of NKG2DL by NKG2D expressing NK cells is more biologically relevant. We investigated the effect of KP reactions in NKG2D signaling within this context. Several computational models of NKG2D signaling(31, 52, 53) have considered KP though the presence of KP in NKG2D signaling has not been validated in experiments. Nevertheless, because of its possible relevance in regulating channel capacity in signaling events in TCR(46), we investigated if the presence of KP in activating NKG2D and inhibitory KIR2DL1 signaling plays any role in affecting the channel capacity or the MI values.

The signaling model investigated until now contains irreversible KP steps for NKG2D and KIR2DL1 signaling. These are indicated in Table S1. We created three different cases where we removed the KP steps (a) from activating NKG2D signaling, (b) from inhibitory KIR2DL1 signaling, and (c) from both. Since, the lifetime of the NKG2DL-NKG2D complex in the model is ∼ 30s, the removal of the KP is likely to have a smaller effect at very early times (≤ 1min) in NKG2D signaling; in contrast, the lifetime of the HLA-C-KIR2DL1 complex is ∼ 0.7s, therefore, removal of KP in KIR2DL1 signaling will have a more noticeable effect in inhibitory signaling. As expected, we found removing KP from NKG2D signaling does not lead to appreciable changes in the mean pVav1 copy numbers (Fig. S5). We have also checked this by solving the rate equations numerically with and without KP for the activating receptors in the disjoint case. However, removing KP from the KIR2DL1 decreased pVav1 copy numbers when KIR2DL1 and NKG2D signaling interact (Fig. S5). We computed MI and CC values for the cases (a-c) for the disjoint, homogeneous, and fully overlapped spatial distributions of NKG2D and KIR2DL1 receptors.

In the case of disjoint clusters for the three different possibilities of turning off KP given above, we found that the values of MI and channel capacity are essentially unaffected and remain greater than unity (**Figures 5A and 5D**). Since the KIR2DL1 receptors are spatially separated from the NKG2D molecules for disjoint clusters removal of KP in the inhibitory reactions has negligible effect on the pVav1 generated via NKG2D signaling; as argued above removal of KP from NKG2D signaling has limited effect on pVav1 production, therefore, the MI and CC values are largely unaffected by removal of KP in the signaling kinetics in the disjoint case.

**Figure 5.**
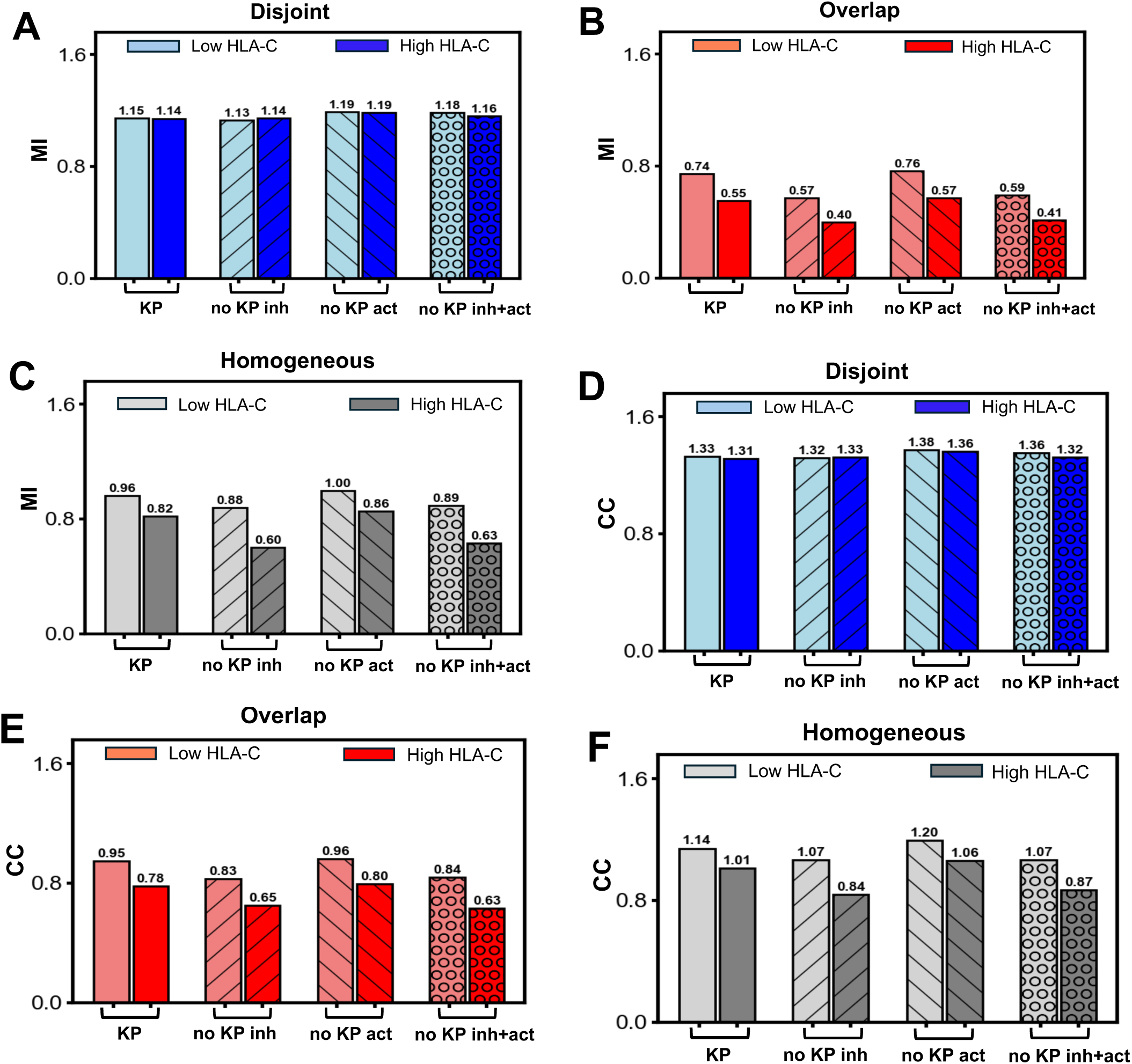
Mutual Information of signal discrimination when kinetic proofreading is removed from activating and inhibitory signaling. Shows the MI values for **(A)** disjoint, **(B)** overlapping, and **(C)** homogeneous cases when KP is removed from NKG2D signaling alone (“no KP act.”), KIR2DL1 signaling alone (“no KP inh.”), or simultaneously from NKG2D and KIR2DL1 signaling (“no KP act.+inh.”). The values when KP is present in both NKG2D and KIR2DL1 (“KP”) are shown as a reference. The values of the CC for **(D)** disjoint, **(E)** overlapping, and **(F)** homogeneous cases are also shown. Note removing KP in the disjoint case or the overlap case leaves the MI values above unity or below unity as they were in the reference case when KP is present (case “KP”). However, removing KP from the inhibitory signaling or both signaling decreases the CC below unity in the homogeneous case for high HLA-C concentration when the CC values for reference case when KP is present in both (“KP”) are above unity.

In the homogeneous and the fully overlapped case, removal of KP from KIR2DL1 signaling decreases MI and CC values (**Figures 5B-C, 5E-F**). One case of interest is in homogeneous case where removal of KP in KIR2DL1 signaling decreases the CC from above unity to less than unity when target cells express high HLA-C abundances (**Figure 5F**). The removal of KP in the inhibitory receptors increases dephosphorylation, since irreversible dissociation of inhibitory complexes is eliminated, thus decreasing pVav1 number and CC. Removing KP from NKG2D signaling alone slightly increases the MI and CC values since the lifetime of complexes that promote phosphorylation do not undergo irreversible dissociation; the time scales, as discussed earlier, are such that no qualitatively relevant changes (such as well below to above unity) occur. When the KP steps are removed simultaneously from NKG2D and KIR2DL1, the changes in the inhibitory signaling dominated over the changes in the activating signaling resulting in lower pVav1 abundances compared to the case when KP is present in both signaling. The corresponding MI and CC values in the homogeneous and fully overlapped case qualitatively follow the behavior when KP is removed from KIR2DL1 alone (**Figures 5B-C, 5E-F**).

Overall, for the different spatial distribution of NKG2D and KIR2DL1 in the resting state, removal of KP makes very little difference. The removal of KP from activating NKG2D and inhibitory KIR2DL1 signaling makes only marginal changes in signal discrimination ability in the disjoint configuration. In the homogeneous cases there are no qualitative differences except in one case (high HLA-C and no KP in the inhibitory sector). In the overlapping case the MI and CC values changes follow expectations for different KP choices but remain well below unity and are not important.

## Discussion

Imaging experiments have demonstrated spatial aggregation of several activating and inhibitory NKRs in the plasma membrane on sub-micron length scales in resting NK cells. We have studied a detailed signaling kinetic model of activating and inhibitory NKRs to evaluate the roles of the observed spatial organization in affecting the ability of NK cells to discriminate between healthy and diseased target cells based on abundances of cognate ligands. The different time scales that arise due to diffusion of cognate ligands and the rates of biochemical signaling reactions determine the early time (< 1 min) signaling behavior. We showed that disjoint clustering of the inhibitory KIR2DL1 and activating NKG2D receptors yield the best response in terms of the ability to the NK cells to discriminate between high and low abundances of activating NKG2DL ligands on short time scales (∼ 20s) regardless of the abundances of the co-expressed inhibitory ligand HLA-C. We quantified this using mutual information measures. This study provides an example of enhancing signal discrimination by local biochemical reactions by exploiting spatial clustering. However, when the clusters of NKG2D and KIR2DL1 fully overlap in the resting state, the NK cells are no longer able to separate low and high concentrations of NKG2D ligands. This inability in separating the abundances of activating ligands can be associated with the inability of NK cells to respond to infected or tumor target cells expressing higher abundances of activating ligands such as ULBP1-6 or MICA/B.

We comment on the potential implications of our results on a different aspect of NK cells. Disjoint or overlapping submicron scale clusters of opposing NKRs can influence NK cell development. Developing NK cells go through an education process where NK cells that do not receive inhibitory signals due to inhibitory NKRs not being expressed or inhibitory NKRs not finding any inhibitory ligand to interact (e.g., NK cells in MHCI KO mice) are rendered hyporesponsive (41). This development program ensures that these NK cells in the periphery are not autoreactive. Guia et al.(30) observed inhibitory Ly49 receptors and activating NKRs (e.g., NK1.1) become confined together in hyporesponsive uneducated NK cells in MHC class I deficient mice where these NK cells generated diminished Ca++ flux when stimulated with NK1.1 antibodies, whereas those NKRs remained in separate compartments in educated NK cells in wild-type mice. These observations are consistent with our prediction that overlapping clusters of activating and inhibitory NKRs in the resting state gives rise to lower pVav1 abundances and potentially to hyporesponsive effector functions in response to stimulation with higher concentrations of activating ligands. Thus, educated NK cells expressing disjoint clusters of opposing NKRs leads to competent discrimination of heathy and diseased target cells expressing low and high concentrations of activating ligands, respectively, whereas the non-educated NK cells are unable to execute this discrimination.

We used mutual information and channel capacity to quantify the ability of NK cells to discern different concentrations and distributions of activating and inhibitory ligands expressed on target cells. The channel capacity (*CC*) provides an upper limit of the number of different signals (∼2*^CC^*) that can be distinguished by the system, and the input distribution of the signal corresponding to the channel capacity provides the optimal distribution of the signal that NK cells discriminate maximally. We show that the input distributions of activating ligand concentrations corresponding to the channel capacity, that yields maximal discrimination is bimodal in nature with peaks at low and high concentrations when the clusters of activating and inhibitory NKRs are disjoint. Similar bimodal distributions are observed in MICA/ULBP3 expressions in HCMV infected human MRC-5 fibroblast target cells(13). However, the expression of NKG2D ligands in tumor cells appeared to follow unimodal distributions(14, 54). Our calculation of mutual information (MI) when the abundances of NKG2DL are distributed in a uniform distribution showed smaller but order of unity magnitude when the clusters of NKRs are disjoint, suggesting the NK cells would still be able to separate target cells with low and high expressions of unimodally distributed NKG2DL. Our calculations also showed if NKRs are distributed homogeneously on the plasma membrane the mutual information becomes slightly smaller than unity when MHC class I expressions are higher (e.g., healthy cells) than that of the tumor cells(**Figure 4B**), however, having NKRs present in disjoint clusters gives rise to a value of mutual information above unity demonstrating a clear discrimination between high and low activating signals in this scenario. This points to the advantage of the formation of submicron scale NKR clusters in the resting state towards increased separation NK cell responses against tumor cells.

The early time NKG2D signaling kinetics is diffusion limited as the average time for an NKG2DL to find an NKG2D receptor is about 10-20s depending on the number of NKG2DL which is much faster (∼10 times) than the time scales of the local binding/unbinding rates of the receptor-ligand pair. This is because of the slower diffusion time scales (∼0.01(μm)^2^/s) of the NKG2DL molecules and low density of NKG2D clusters on target cells (∼ 4 non-empty clusters/(μm)^2^). The diffusion limitation in the receptor ligand complex formation produces a slower increase in pVav1 abundances for moderate NKG2DL concentrations (∼ 15 molecules/(μm)^2^) in the model which starts to saturate after 20sec. The kinetics of the MI values show that MI values greater than unity are reached in about 10secs (see **Figure 4C**) for disjoint NKR clusters demonstrating that the NK cell achieves the discrimination between high and low NKG2DL abundances on short time scales. This could be particularly relevant as NK cells lysis of target cells by NK cells through perforin/granzyme B can occur within minutes after stimulation(55, 56).

The absence of kinetic proofreading in NKG2D or KIR2DL1 biochemical signaling kinetics did not appreciably affect the values of MI or CC for NKRs residing in disjoint or overlapping. This suggests that KP in the membrane proximal signaling kinetics of NKRs, unlike T cell receptor signaling(46), does not play a critical role in signal discrimination for NK cells. This is because NK cells need to discriminate between healthy and diseased target cells expressing different abundances of high affinity ligands cognate to activating and inhibitory NKRs; on the other hand, T cells require to sense the presence of few high affinity ligands among a large abundance of low affinity self-ligands. The KP model proposed by McKeithan, Hopfield and Ninio achieves a sensitive separation between ligands that form receptor-ligand complexes with slightly different half-lives, however, for a fixed affinity, the concentrations of the active complex in the KP model increases almost linearly and then reaches saturation with ligand concentrations – this is particularly relevant for different agonist and self-ligands of TCRs. However, the ligands cognate to NKG2D and KIR2DL1 are of higher affinity than typical ligands for the TCR, and NK cells need to discriminate between high and low abundances of these ligands in healthy and tumor cells, thus, the presence of KP is does not affect signal discrimination due to variation of ligand concentration.

NKG2D stimulation needs to be combined with stimulation by another activating NKR (e.g., 2B4) to induce NK cell cytotoxicity(57). Similar pairwise synergy involving several activating NKRs and adhesion receptors have been reported for NK cells(57). The joint stimulation by NKG2D and 2B4 induces a dual phosphorylation of a pair of tyrosine residues in an adaptor protein SLP76(38) which upon the tyrosine phosphorylation binds to Vav1. Therefore, the discrimination of healthy and target cells based on NKG2DL concentrations are likely to be influenced by target cell expressions of ligands for 2B4. This would be an interesting direction to use our framework to investigate the role of pairwise synergy between activating receptors in affecting ligand discrimination by NK cells. Galstyan et al.(58) showed how enzymatic reactions can exploit spatial concentration gradients of substrates to proofread signals. In our work we have examined how NK cells can exploit spatial organization of membrane receptors, by clustering, to improve discrimination between high and low signal amplitudes. This points to how cells can use local biochemical reactions combined with spatial diffusion to tune time scales or enhance biological function.

## Materials and Methods

### Spatially resolved stochastic simulations

We simulate spatiotemporal signaling kinetics in a quasi-two-dimensional simulation box of area 3.6 × 3.6 μm^2^. The simulation box represents (i) the region of contact between the cognate ligands expressed on the target cell and the NKRs bound to the NK cell surface, and (ii) the transmembrane region of the NK cell where early time signaling reactions post-ligand-receptor binding take place. The simulation box is divided into three dimensional chambers of volume of *l*_0_ × *l*_0_ × *z* (nm), where *l*_0_=40nm, and different *z* values are chosen depending on the location of the protein species. The molecules are assumed to be well-mixed within the chambers. We use a software package SPPARKS (https://spparks.github.io/) to perform the stochastic simulation using Gibson-Bruck implementation of the Gillespie algorithm. The following biochemical reactions and the diffusion moves are used in the model.

#### Biochemical reactions

The model includes plasma membrane bound NKG2D and KIR2DL1 receptors and the kinase SFK, and the cytosolic Vav1 and SHP1 molecules in NK cells. The ligands NKG2DL and HLA-C reside in the plasma membrane of the target cell. The molecules of the above protein species in the chambers interact following biochemical reactions described in Table S1. The propensities given there are obtained as follows. We set *z*=*l*_0_ for protein species that reside in the plasma membrane, and set *z*=1.0 μm for species that reside in the cytosol. The second order rate (k_on_) for binding of two molecules (e.g., NKG2D binding NKG2DL) are provided in the literature in units of (density)^-1^s^-1^ (e.g., (μM)^-1^s^-1^). We converted these rates into propensities which defines the inverse timescale at which a particular reaction occurs. The binding rate k_on_ is converted into the propensity ∝ k_on_/ν, by dividing the rate k_on_ by a volume factor, ν defining the volume of the region where the well-mixed reaction takes place. The binding of NKR and their cognate ligands take place when they are interacting domains are separated by d_recep-lig_ ∼ 2nm, thus ν_recep-lig_=*l*_0_×*l*_0_×*d*_recep-lig_ for the receptor ligand binding reactions. The binding of SFK to NKR-ligand species occur in the plasma membrane, thus we set ν_recep-SFK_ =*l*_0_×*l*_0_× *l*_0_. The volume factor, ν_recep-Vav1/SHP1_, binding of cytosolic proteins (e.g., Vav1 or SHP1) to the transmembrane domains of the receptors is set to ν_recep-Vav1/SHP1_*= l*_0_×*l*_0_×z.

#### Spatial movements

We considered diffusive movements of free molecules residing in the plasma membrane and the cytosol. We did not include any cytoskeletal forces or movements of receptors due to changes in the apposing NK- and target cell plasma membranes at the immunological synapse as we considered early time (< 1 min) kinetics. The diffusive movements are modeled as hops of molecules from a chamber to one of its four nearest neighboring chambers with a rate *D*/*l*_0_^2^, where D is the diffusion constant of the molecule. For the disjoint and overlapping cases, we restrict diffusion of the NKRs such that the NKRs residing in a cluster (or outside the cluster) do not hop to out the cluster (or in the cluster). The unattached molecules of SFK and SHP1, and Vav1 freely diffuse throughout the simulation box according to their respective diffusion coefficients.

The reactions, their rates and propensities, the diffusion rates, the initial concentrations of the molecules, and the relevant references are shown in Table S1.

#### Initial configuration

##### Formation of submicron clusters

The submicron clusters of NKG2D and KIR2DL1 are created by choosing the clusters uniformly and randomly from the images and placing those in the simulation box with a uniform distribution where we restrict any overlap between the clusters of the same type. A chamber falling within or outside an NKR cluster is marked 1 or 0, respectively. The NKRs diffuse within the chambers marked 1 or 0, but do not hop from a chamber marked 1 to 0 or vice versa. The numbers of the clusters in the simulation box are estimated by keeping the densities of the respective clusters equal to that of those in the binary maps of the superresolution images in refs (16) (17). To generate the initial configurations for non-overlapping or disjoint NKG2D and KIR2DL1 clusters, the NKG2D clusters are nucleated first in the simulation box, then the KIR2DL1 clusters are nucleated with the restriction that the NKG2D and KIR2DL1 clusters cannot overlap. For the initial configurations with fully overlapping NKG2D and KIR2DL1 clusters, the NKG2D clusters are nucleated first, then the center of mass of the KIR2DL1 are made to coincide with that of the NKG2D clusters. Since the density of KIR2DL1 clusters are higher, once all the NKG2D clusters are overlapped with the KIR2DL1 cluster, the rest are distributed randomly with a restriction of not overlapping with the NKG2D clusters. The NKG2D and KIR2DL1 molecules are placed randomly in their respective clusters. If a cluster does not acquire any NKR, we remove it from the simulation. The fraction of deleted clusters is 56.4% for NKG2D and 0% for KIR2DL1. <u>Homogeneous distribution:</u> NKG2D and KIR2DL1 are distributed uniformly and randomly. Since there are no clusters, all the chambers are marked 0.

The proteins SFK, Vav1, and SHP1 are distributed homogeneously throughout the simulation box.

#### Boundary conditions

We use periodic boundary conditions in the x- and y- directions, and impenetrable boundary conditions along the z- directions.

#### Calculation of the two-point correlation function

We computed the two-point correlation function *C*(*r*) for the binary maps of the superresolution images. The binary maps assign a binary variable *S_NKR_*(*x_i_*,*y_i_*) equal to 1 or 0 at a spatial location (*x_i_*, *y_i_*) if its occupied or not occupied by an NKR. The extracted binary variables are placed on a square lattice with a total of *N_total_* sites with a lattice constant=1. We computed the mean (*µ_NKR_*), variance (*σ_NKR_*), and the two-point correlation function (*C_NKR_*(*r*)) using *S_NKR_*(*x_i_*, *y_i_*) as defined below:

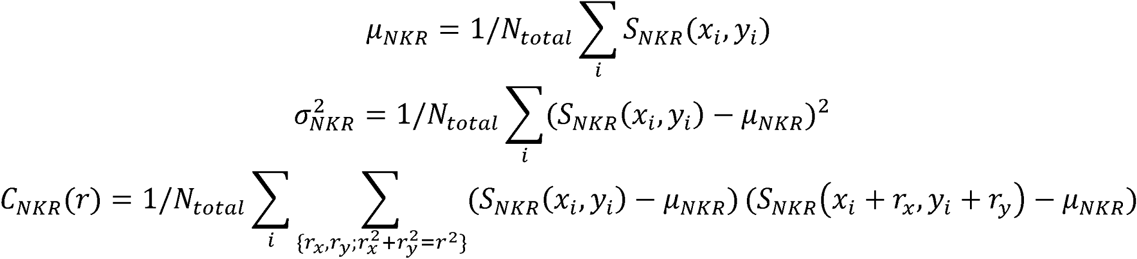

The summation over *r_X_* and *r_Y_* indicates average over all neighbors that reside at a distance *r* from (*x_i_*,*y_i_*. We compare the variance scaled two point correlation function, 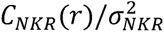, in Fig. 2 for the clusters obtained from the superresolution images and the random organization of those clusters.

##### Extraction of image data

The initial NKG2D and KIR2DL1 clusters were extracted from the binary maps of the superresolution images available in Fig. 1B in (16) in 5× 5 *µm* region, and in Fig. 4A (top right panel) in (17) in 2×2 *µm* region, respectively. Signal intensities of the original image were extracted in pixels of 40nm×40 nm, and the intensities ≥ 100 (or <1000) were set to 1 or occupied (or, unoccupied or 0).

##### Calculation of the MI and CC

One can show that *MI*(*o*;*s*) - *H*(*s*) - *H*(*s*|*o*), the difference between the Shannon entropy of the signal, *H*(*s*) - - ∑*_s_ Q*(*s*)*Log*_2_*Q*(*s*), and the conditional entropy defined by the conditional probability distribution, 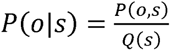 *H*(*s*|*o*) − Σ_*o*_ Σ_*s*_ *P*(*o,s*)*Log*_2_*P*(*s*|*o*). Since *H*(*s*) can be viewed as the uncertainty in the signal while *H*(*s*|*o*) is the uncertainty in the signal given the output, the mutual information is the uncertainty in the signal reduced by the information contained in the output. Thus, larger the value of the MI, the greater is the fidelity of the output to the input. In fact, *MI*(*o*; *s*) is the Kullback-Leibler divergence between *P*(*o*, *s*) and the product distribution *R*(*a*)*Q*(*s*). The MI depends on the input distribution Q(s) and the capacity is defined as the maximum MI as a function of Q. It is not easy to perform this maximization since P(o|s) itself is a function of Q, in addition to it appearing explicitly. The Blahut-Arimoto algorithm overcomes this by introducing an auxiliary function Φ(o|s) that represents an arbitrary set of conditional probabilities and the function 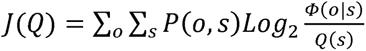 is maximized iteratively by finding Q(s) for a given Φ and given Φ, finding a new Q. The procedure has been shown to converge with an explicit criterion for stopping given a required accuracy and is easy to implement numerically. We computed MI and CC for seven input values of s. Increasing the number of inputs changes the magnitudes of MI and CC changes by a small (∼1%) amount for most of the cases (Fig. S3), therefore, we report the results for seven inputs.

## Data and code availability

The SPPARKS codes to run the simulations are available at our GitHub page https://github.com/drsamalik/NKG2D_KIR2DL1_clustering.

## Acknowledgement

This work was supported by the NIH Office of the Director award R01-AI 146581 to Jayajit Das. We also acknowledge funding from the Nationwide Children’s Hospital and the Ohio Supercomputing Center for the use of MATLAB.

## Author Contribution

Saeed Ahmad: Formal analysis; investigation; methodology; software; validation; visualization; writing – original draft; writing – review and editing. Debangana Mukhopadhyay: Formal analysis; investigation; methodology; software; validation; writing – original draft; writing – review and editing. Rajdeep Grewal: Formal analysis; investigation; methodology; Ciriyam Jayaprakash: Conceptualization; investigation; methodology; project administration; supervision; writing – original draft; writing – review and editing. Jayajit Das: Conceptualization; funding acquisition; investigation; methodology; project administration; supervision; writing – original draft; writing – review and editing.

## Notes

### Competing Interest Statement

The authors have declared no competing interest.

