## Supplementary Material for "Spatial statistics of submicron size clusters of activating and inhibitory Natural Killer cell receptors in the resting state regulate early time signal discrimination"

**Supplementary Text1: Calculation of the probability that a site occupied by NKG2D does not contain any KIR2DL1 molecule**

Consider an area of L× L in the plasma membrane of an NK cell. Experiments show that KIR2DL1 molecules suppress NKG2D signaling the most when NKG2D and KIR2DL1 are 40nm apart [1]. The area is divided into N*_l_*, number of small chambers of size *l*_0_× *l*_0_ where *l*_0_=40nm. If a particular small chamber is occupied by NKG2D, the probability that a KIR molecule will occupy that chamber when KIRs are distributed from uniform random distribution is *p*=1/N*_l_* = *l*_0_^2^/L^2^. Therefore, the probability that the chamber is not occupied by a KIR is (1-*p*). If we draw N number of KIR molecules, then the probability that the chamber is not occupied by any KIR is then q=(1-p)^N^=(1- *l*_0_^2^/L^2^)^N^ =exp(N*ln*(1- *l*_0_^2^/L^2^)) ≈ exp(-*l*_0_^2^/ξ^2^) where ξ=L/√N ≡ average distance between the KIRs. For a ξ=34nm which is the average distance between KIRs distributed homogeneously in the membrane of a primary NK cell, ~25% of the NKG2D molecules will not be partnered by any KIR.

**Supplementary Text2: Calculation of Kullback-Leibler divergence between pVav1 distributions**

The pVav1 distributions shown in Figure 3 (E-G) and Fig. S1 (F-N) can be fit by the Gamma distribution given by $\gamma\left( x \right)=\frac{b^{a}}{\Gamma(a)}x^{a-1}e^{-bx}$. The parameters a and b for different NKG2DL numbers, for high and low HLA1-C, and for all three spatial distributions of receptors can be found in GitHub. This permits another quantitative characterization of how well low and high NKG2DL values can be differentiated in the three scenarios.

The Kullback-Leibler divergence or the relative entropy $D_{KL}(P||Q)$ is a measure of how much any probability distribution Q differs from a given probability distribution P and is defined to be $\int dx p\left( x \right) log(\frac{p\left( x \right)}{q\left( x \right)})$ where p and q are the corresponding densities. An analytic expression for the K-L divergence when	$p \sim\gamma(a_{1},b_{1})$ and	$q\sim\gamma(a_{2},b_{2})$ can be calculated and is given by

$$KL(a_{1},b_{1} ||a_{2},b_{2})= a_{2}Log[b_{1}/b_{2}] + Log[\Gamma(a_{2})/\Gamma(a_{1})] + (a_{1}-a_{2}) \psi(a_{1}) + \left( b_{2}-b_{1} \right)a_{1}/b_{1}$$

Given the fitting parameters we find that the KL divergence between the pVav1 distributions for NKG2DL = 400 and 20 with low HLA-C are 11.1, 2.2, and 4.7 for the disjoint, overlapping, and homogeneous cases, respectively. This is another measure of how much better the disjoint cluster configuration performs in discriminating between high and low ligand concentrations. Even for closer values of NKG2DL of 200 and 40 the KL divergence is 7.2 for the disjoint case (larger than in the other two scenarios when the NKG2DL numbers are closer) and 2.7 and 4.3 for the overlapping and homogeneous cases.

I

**Figure S1. Characterizing the variations of the response pVav1, as a function of the strength of the input signal, NKG2DL in the presence of HLA-C, for disjoint, homogeneous, and overlapping distributions of receptors. (A)** The mean value of the pVav1 distribution as function of the NKG2DL number) at 20s, when the HLA-C number is high (=11686). The behavior for disjoint clusters is shown in blue, for overlapping clusters in red and the homogeneous case in grey. The mean pVav1 number is observed to be highest for disjoint clusters, followed by homogeneous and overlapping cases. The variance indicated by the vertical bars; the responses to 100 and 400 NKG2DL do not overlap for the disjoint clusters in contrast to the overlapping case. **(B)-(C)** The mean pVav1 as a function of NKG2DL at 40s or low (B) and high(C) HLA-C values of 4213 and 11686, respectively. The response is diminished when the HLA-C number increases in the homogeneous case in contrast to the disjoint case. The response is much weaker in the overlapping case both here and at 20s in (A). **(D)-(E)** The temporal evolution of mean pVav1 is shown for the three cases with HLA-C=4213 for NKG2DL=40 in (D) and 200 in (E) respectively. The time scale for saturation is set by diffusion that determines the rate at which the cognate ligands find and bind to the NKG2D receptors (53 in our simulation). A higher concentration of NKG2DL results in a more rapid approach to steady state and higher values of pVav1 at any time. **(F)-(N)** pVav1 distributions are displayed for different NKG2DL numbers (20, 40, 100, 150, 200, 400) for the three different spatial distribution of receptors at 20s for high HLA-C (F-H) and at 40s for low HLA-C (I-K) and high HLA-C (L-N). The distributions at low and high NKG2DL are well-separated for disjoint clusters allowing better discrimination between them and leading to higher mutual information MI values as shown in Figure 4 in the main text. In the case of overlapping clusters, the distributions are poorly separated. For the homogeneous case, the results are closer to the disjoint case though less well-separated. There is significant overlap of the distributions for NKG2DL = 200, 300, and 400 implying that the system cannot discriminate between them.

**Figure S2. Mutual Information and Channel capacity at t=40s.** Bar Charts showing values of Mutual Information (MI) in (A) and Channel Capacity (CC) in (B) for the three conditions—disjoint, homogeneous, and overlap—at time 40s. Light shading represents low HLA-C values (4213), while dark shading represents high HLA-C numbers (11686). Disjoint clusters lead to the highest values of MI and CC, followed by homogeneous and overlapping case conditions. The decrease in the values when HLA-C number is increased reflects the increased number of KIR-HLA-C complexes and its descendants that inhibit pVav1 production.

**Figure S3**. **Variation of MI and CC with different numbers of inputs used in evaluating MI and CC.** (A) MI and (B) CC values evaluated for different number of inputs for disjoint, overlapping, and homogeneous cases. The maximum percentage changes as the number of inputs is increased from 7 are shown on the panels.

**Figure S4. Variation of the mean total number of NKG2DL bound complexes with NKG2DL concentrations.** The mean number of the sum of all the NKG2DL bound complexes increase first and then saturate with increasing concentrations of NKG2DL. The number of (=53) available NKG2D receptors sets the maximum value of the mean. At lower concentrations of NKG2DL (e.g., 100), the mean value increases more for the time interval t=20s to t=40s, as compared to t=40s to t=60s, indicating the diffusion limitation in the formation of the complexes as NKG2DL molecules diffuse to find NKG2D receptors within the clusters.

**Comparison of mean pVav1, MI and CC for disjoint case, low HLA-C:**

**Figure S5**. **Kinetics of mean pVav1 numbers for the disjoint and the overlapping cases in the absence of KP.** **(A-C)** Shows the kinetics for the disjoint case, where the mean pVav1 shown in (A-B) remains largely unchanged between when the KP is absent or present (no KP). The SHP1 as shown in (C) increases in the absence of KP, however, since the inhibitory KIR2DL1 clusters are disjoint the increase does not affect mean pVav1. **(D-F)** Kinetics for the overlapping case, where the mean pVav1 shown in (D-E) decreases under the where the KP is absent. The lower pVav1 is due to the increased inhibition by the higher number of bound SHP1 molecules shown in (F) when KP is absent compared to the case when KP is present.

**Table S1**

A. **List of reactions and parameter values used in the model:**

k_on_ denotes binding rate for species *x* and *y*. k_off_ denotes unbinding rate of complex *x*-*y*. k_cat_ is the catalytic rate of phosphorylation/dephosphorylation of *x* by enzyme. The parameter values are taken from Ref. [2].

| ***Activating NKG2D receptor reactions (for all cases)*** | | | | | |
| --- | --- | --- | --- | --- | --- |
| **Comment** | **Reactions** | **k_on_ (s^-1^)** | | **k_off_(s^-1^)** | **k_cat_(s^-1^)** |
| NKG2D binding/unbinding with NKG2DL | NKG2D + NKG2DL ⇌ NKG2D-NKG2DL | 12.385 | | 0.023 |  |
| Binding/unbinding of SFK with DAP10 | NKG2D-NKG2DL + SFK ⇌ NKG2D-NKG2DL-SFK | 109.370  (50) | | 0.006 |  |
| Phosphorylation of tyrosine residues via SFK | NKG2D-NKG2DL-SFK → pNKG2D-NKG2DL + SFK |  | |  | 2.189 |
| Vav1 binding/unbinding with pITAM | pNKG2D-NKG2DL + Vav1 ⇌ pNKG2D-NKG2DL-Vav1 | 0.634 | | 0.01 |  |
| Binding/unbinding of SFK with pITAM-Vav1 | pNKG2D-NKG2DL-Vav1 + SFK ⇌ pNKG2D-NKG2DL-Vav1-SFK | 781.172  (50) | | 0.028 |  |
| Phosphorylation of Vav1 by SFK | pNKG2D-NKG2DL-Vav1-SFK → pNKG2D-NKG2DL-pVav1 + SFK |  | |  | 0.776 |
| Dephosphorylation of pVav1 | pNKG2D-NKG2DL-pVav1 → pNKG2D-NKG2DL-Vav1 |  | |  | 1.0 |
| Dephosphorylation of pDAP10 | pNKG2D-NKG2DL → NKG2D-NKG2DL |  | |  | 2.0 |
| Dissociation of free pVav1 | pNKG2D-NKG2DL-pVav1 ⇌ pNKG2D-NKG2DL + pVav1 | 0.634 | | 0.01 |  |
| Dephosphorylation of free pVav1 | pVav1 → Vav1 |  | |  | 1.0 |
| ***Inhibitory receptor KIR2DL1 reactions (for all cases)*** | | | | | |
| **Comment** | **Reactions** | **k_on_ (s^-1^)** | | **k_off_ (s^-1^)** | **k_cat_ (s^-1^)** |
| Binding/unbinding of KIR and HLA | KIR + HLA ⇌ KIR-HLA | 50.0 | | 1.0 |  |
| Binding/unbinding of SFK | KIR-HLA + SFK ⇌ KIR-HLA-SFK | 50.0 | | 0.006 |  |
| phosphorylation of ITIM | KIR-HLA-SFK → pKIR-HLA + SFK |  | |  | 2.189 |
| Binding/unbinding of SHP1 | pKIR-HLA + SHP1 ⇌ pKIR-HLA-SHP1 | 0.375 | | 0.0005 |  |
| Binding/unbinding of KIR and NKG2D complexes | pKIR-HLA-SHP1 + pNKG2D-NKG2DL-pVav1 ⇌ pKIR-HLA-SHP1-pNKG2D-NKG2DL-pVav1 | 158 (50) | | 0.01 |  |
| Dephosphorylation of ITIM | pKIR-HLA → KIR-HLA |  | |  | 2.0 |
| Dephosphorylation of Vav1 | pKIR-HLA-SHP1-pNKG2D-NKG2DL-pVav1 → pKIR-HLA-SHP1 + pNKG2D-NKG2DL-Vav1 |  | |  | 10.0 |
|  | pKIR-HLA-SHP1 + pVav1 ⇌ pKIR-HLA-SHP1-pVav1 | 6.342 | | 0.01 |  |
| Dephosphorylation of Vav1 | pKIR-HLA-SHP1-pVav1 → pKIR-HLA-SHP1 + Vav1 |  | |  | 10.0 |
| ***Reactions for implementing kinetic proofreading (KP)*** | | | | | |
| **Comment** | **Reactions** | **k_on_ (s^-1^)** | | **k_off_ (s^-1^)** | **k_cat_ (s^-1^)** |
| NKG2DL unbinding | NKG2D-NKG2DL-SFK → NKG2D + NKG2DL + SFK |  | | 0.023 |  |
|  | pNKG2D-NKG2DL-Vav1 → NKG2D + NKG2DL + Vav1 |  | | 0.023 |  |
|  | pNKG2D-NKG2DL → NKG2D + NKG2DL |  | | 0.023 |  |
|  | pNKG2D-NKG2DL-Vav1-SFK → NKG2D + NKG2DL + Vav1 + SFK |  | | 0.023 |  |
|  | pNKG2D-NKG2DL-pVav1 → NKG2D + NKG2DL + pVav1 |  | | 0.023 |  |
|  | pKIR-HLA-SHP1-pNKG2D-NKG2DL-pVav1 →  pKIR-HLA-SHP1 + NKG2D + NKG2DL + Vav1 |  | | 0.023 |  |
| HLA unbinding | pKIR-HLA → KIR + HLA |  | | 1.0 |  |
|  | KIR-HLA-SFK → KIR + HLA + SFK |  | | 1.0 |  |
|  | pKIR-HLA-SHP1 → KIR + HLA + SHP1 |  | | 1.0 |  |
|  | pKIR-HLA-SHP1-pVav1 → KIR + HLA + SHP1 + Vav1 |  | | 1.0 |  |
|  | pKIR-HLA-SHP1-pNKG2D-NKG2DL-pVav1 → KIR + HLA + SHP1 + pNKG2D-NKG2DL-pVav1 |  | | 1.0 |  |
| ***Reactions when KP is absent in NKG2D signaling*** | | | | | |
| **Comment** | **Reactions** | **k_on_ (s^-1^)** | **k_off_ (s^-1^)** | | **k_cat_ (s^-1^)** |
| NKG2DL unbinding | NKG2DL + NKG2D-SFK ⇌ NKG2D-NKG2DL-SFK | 12.38 | 0.023 | |  |
|  | NKG2DL + pNKG2D-Vav1 ⇌ pNKG2D-NKG2DL-Vav1 | 12.38 | 0.023 | |  |
|  | NKG2DL + pNKG2D ⇌ pNKG2D-NKG2DL | 12.38 | 0.023 | |  |
|  | NKG2DL + pNKG2D-Vav1-SFK ⇌ pNKG2D-NKG2DL-Vav1-SFK | 12.38 | 0.023 | |  |
|  | NKG2DL + pNKG2D-pVav1 ⇌ pNKG2D-NKG2DL-pVav1 | 12.38 | 0.023 | |  |
|  | NKG2DL + pKIR-HLA-SHP1-pNKG2D-pVav1 ⇌ pKIR-HLA-SHP1- pNKG2D-NKG2DL-pVav1 | 12.38 | 0.023 | |  |
|  | NKG2DL + pKIR-SHP1-pNKG2D-pVav1 ⇌ pKIR-SHP1-pNKG2D-NKG2DL-pVav1 | 12.38 | 0.023 | |  |
| Dissociation of complexes | NKG2D-SFK → NKG2D + SFK |  | 0.006 | |  |
|  | pNKG2D-Vav1 → pNKG2D + Vav1 |  | 0.01 | |  |
|  | pNKG2D-Vav1-SFK → pNKG2D-Vav1 + SFK |  | 0.006 | |  |
|  | pKIR-SHP1-pNKG2D-Vav1 → pKIR-SHP1 + pNKG2D-Vav1 |  | 0.01 | |  |
|  | pKIR-SHP1-pNKG2D-pVav1 → pKIR-SHP1 + pNKG2D-pVav1 |  | 0.01 | |  |
|  | pNKG2D-pVav1 → pNKG2D + pVav1 |  | 0.01 | |  |
| Phosphorylation of Vav1 by SFK | pNKG2D-Vav1-SFK → pNKG2D-pVav1 + SFK |  | 0.776 | |  |
| ***Reactions when KP is absent in KIR2DL1 signaling*** | | | | | |
| **Comment** | **Reactions** | **k_on_ (s^-1^)** | **k_off_ (s^-1^)** | | **k_cat_ (s^-1^)** |
| KIR unbinding | HLA + pKIR ⇌ pKIR-HLA | 50.0 | 1.0 | |  |
|  | HLA + pKIR-SHP1 ⇌ pKIR-HLA-SHP1 | 50.0 | 1.0 | |  |
|  | HLA + pKIR-SHP1-pNKG2D-NKG2DL-pVav1 ⇌ pKIR-HLA-SHP1- pNKG2D-NKG2DL-pVav1 | 50.0 | 1.0 | |  |
|  | HLA + pKIR-SHP1-pNKG2D-pVav1 ⇌ pKIR-HLA-SHP1-pNKG2D-pVav1 | 50.0 | 1.0 | |  |
|  | HLA + KIR-SFK ⇌ HLA-KIR-SFK | 50.0 | 1.0 | |  |
| Dissociation of complexes | pKIR-SHP1 → pKIR + SHP1 |  | 0.0005 | |  |
|  | KIR-SFK → KIR + SFK |  | 0.006 | |  |
| Phosporyation of ITIM | KIR-SFK → pKIR + SFK |  |  | | 2.189 |
| Dephosphorylation of pVav1 | pKIR-SHP1-pNKG2D-NKG2DL-pVav1 → pKIR-SHP1 + pNKG2D-NKG2DL-Vav1 |  |  | | 1.0 |
|  | pKIR-HLA-SHP1-pNKG2D-pVav1 → pKIR-HLA-SHP1 + pNKG2D-Vav1 |  |  | | 1.0 |
|  | pKIR-SHP1-pNKG2D-pVav1→ pKIR-SHP1+pNKG2D -Vav1 |  |  | | 1.0 |
|  | pNKG2D-pVav1 → pNKG2D-Vav1 |  |  | | 1.0 |
| Dephosphorylation of pDAP10 | pNKG2D → NKG2D |  |  | | 2.0 |
| Dephosphorylation of ITIM | pKIR → KIR |  |  | | 2.0 |

**B. List of protein concentrations used in the model:**

| **Molecule type** | **Number of molecules** | **Concentration** | **Comment** |
| --- | --- | --- | --- |
| KIR2DL1 | 11686 | 882 /µm^2^ | Assuming the diameter of an NK cell to be 10µm and 277,215 molecules /cell reported by ref. [3] |
| NKG2D | 53 | 4/µm^2^ | Number NKG2D are ranging 662-1321 molecules /cell with mean = 995 molecules /cell (ref.[2]) |
| SFK | 32064 | 2420/µm^2^ | 553000 – 1245000 molecules/cell with mean = 760,000 molecules /cell  Concentration = 292,890 to 665,220 nM (402,520 nM = mean) (ref.[2]) |
| Vav1 | 18351 | 1385/µm^2^ | 79,000 to 359,000 molecules/cell  Concentration = 920 – 4200 nM (2300 nM= mean value)  Number of Vav1 per µm^3^ = 554 - 2529 (1385 molecules/ µm^3^) (ref.[2]) |
| SHP1 | 49871 | 3764/µm^2^ | 405,000 to 648,000 molecules/cell  Concentration = 4730 – 7570 nM (6250 nM = mean value)  Number of SHP1 per µm^3^ = 2899 - 4559 (3764 molecules/ µm^3^) (ref.[2]) |
| HLA-C | 4213(low) 11686(high) | 318 /µm^2^ (low)  882/µm^2^ (high) | 100,000 molecules/cell (ref.[2]) |

1.	Toledo, E., et al., *Molecular-scale spatio-chemical control of the activating-inhibitory signal integration in NK cells.* Science advances, 2021. **7**(24): p. eabc1640.

2.	Grewal, R.K. and J. Das, *Spatially resolved in silico modeling of NKG2D signaling kinetics suggests a key role of NKG2D and Vav1 Co-clustering in generating natural killer cell activation.* PLOS Computational Biology, 2022. **18**(5): p. e1010114.

3.	Watzl, C. and D. Urlaub, *Molecular mechanisms of natural killer cell regulation.* Front Biosci, 2012. **17**: p. 1418-32.
